# Comparison of Different Kits for SARS-CoV-2 RNA Extraction Marketed in Brazil

**DOI:** 10.1101/2020.05.29.122358

**Authors:** Ana Karolina Antunes Eisen, Meriane Demoliner, Juliana Schons Gularte, Alana Witt Hansen, Karoline Schallenberger, Larissa Mallmann, Bruna Saraiva Hermann, Fágner Henrique Heldt, Paula Rodrigues de Almeida, Juliane Deise Fleck, Fernando Rosado Spilki

## Abstract

December 2019 marked the begining of the greatest pandemic since Spanish Flu, the disease named Covid-19 that cause severe pneumonia. Until May 19, 2020 more than 4 million and 700 thousand cases were oficially notified with about 316 thousand deaths. Etiological agent of the disease was identified as being a new coronavirus, *Severe acute respiratory syndrome-related coronavirus* (SARS-CoV-2). In this study we compared four different manual methods for RNA isolation and purification for detection of SARS-CoV-2 through qRT-PCR, as well as the extraction quality itself through detection of RNAse P. Magnetic beads-based (MagMax™) and silica column-based (Biopur^®^) methods presented the better performances. Concerning to the mean delay in CT values when compared to MagMax™, TRIzol™, Biopur^®^ and EasyExtract presented 0,39, 0,95 and 5,23 respectively. Agreement between positive and negative results of different methods when compared with the one with better performance MagMax™ was 94,44% for silica column-based method (Biopur®), 88,89% for phenol-chroloform-based method (TRIzol™) and 77,78% for EasyExtract. We aimed to evaluate how reliable each method is for diagnostic purposes and to propose alternatives when usual methods are not available. In this regard, magnectic beads and silica column-based methods are convenient and reliable choices and phenol-chloroform-based method could also be chosen as an alternative.

## INTRODUCTION

December 2019 marked the begining of the greatest pandemic since Spanish Flu, when the first cases of Covid-19 were identified, an atypical pneumonia of unknown origin. Infection origin was initially associated with a seafood market in Wuhan city, Hubei province, China [1–3]. Disease was named as Covid-19 by the World Health Organization (WHO) in a report that was published in February 22, 2020 [4]. The infection had spread rapidly through the Europe and the Americas since the first imported cases were arrived, until May 19, 2020 more than 4 million and 700 thousand cases were oficially notified with about 316 thousand deaths distributed through 216 countries [5].

Etiological agent of the disease was identified as being a new coronavirus originating from bats with a possible relation to a coronavirus identified in pangolins of *Manis javanica* species [2, 3, 6]. The new coronavirus was classified in the same species of the first SARS of 2003 as *Severe acute respiratory syndrome-related coronavirus* (SARS-CoV-2), within the subgenre *Sarbecovirus,* genre *Betacoronavirus,* subfamily *Orthocoronavirinae* and *Coronaviridae* family. Members of this viral Family are mainly spherical and have lipid envelopes with prominent proteins named Spikes, which ones give the crown appearance that inspired family name. Genome is composed by a positive single stranded RNA that measures about 26 - 31 kb and is readed through 6 ORFs [7]. Besides SARS-CoV-2, six other coronavirus are already known by cause infection in humans, among them the SARS and MERS-CoV pandemic viruses that may cause severe disease and pneumonia and the other four human coronavirus (HCoV) that are causal agents of comon colds OC43, 229E, HKU1 and NL63 [8,9].

Covid-19 is a mainly respiratory disease that have as most common sympstoms the fewer, cough, shortness of breath, fadigue and difficulty breathing, other symptoms as chills, sore throat, mialgia, loss of smell and taste may also occur with minor frequency, besides gastrointestinal manifestations as diarrhea, nausea and vomiting. In most severe cases the disease course with severe pneumonia that usually lead to hospitalization and the need to use mechanical respirators [10–12].

Two essential measures are required to control of the pandemic while no vaccines are available: social distancing and large scale testing of the population. Due to high infected number, even testing only symptomatic pacientes who looked for medical care there is still a huge sample volum to do the diagnostic. Thereby, alternative methods may be required when lack of kits and reagents would occur due to high demand, mainly in nucleic acids extraction prior to detection by quantitative reverse transcriptase polymerase chain reaction (qRT-PCR). Here we compared four different manual methods for SARS-CoV-2 RNA isolation and purification aiming detection by qRT-PCR marketed as commercial kits in Brazil.

## MATERIALS AND METHODS

Methods compared in this study include magnetic beads method MagMax™ CORE Nucleic Acid Purificatioin Kit (ThermoFisher™), which was adapted for a manual procedure using magnetic racks, column-based method Mini Spin Plus Kit (Biopur^®^) for total nucleic acid extraction, TRIzol™ (Invitrogen™), a phenol-chloroform-based method for high quality RNA isolation and a simple fast methodology for sample preparation prior to the PCR amplification named EasyExtract (Calpro AS™). With exception of MagMax™ in which an adaption for the manual procedure, the other methods were performed exactly as described by their respectives manufacturers.

### Samples and controls

To compare quality of extraction methods, an inactivated Brazilian reference strain cultivated in VERO-Slam cells gently provided by Dr. Edison Durigon (Universidade de São Paulo) was used as a control, and tested pure and diluted with a dilution factor of 10, eight times into cell culture minimum essential medium (MEM), each dilution corresponded to a final sample in the performance assay. In addition to control, a set of SARS-CoV-2 positive nine clinical samples already tested in our lab was chosen based on cycle threshold value (CT), being three of each CT range in qRT-PCR for SARS-CoV-2 (alloted as low, medium and high).

In order to properly compare the different methods, all final samples were aliquoted separately for each technique and freezed (ultrafreezer −80°C) only one time before total nucleic acids/RNA extractions were done, the same attention was taken with the nucleic acids extracted before analysis by qRT-PCR.

### Manual nucleic acid purification by magnetic beads

As mentioned above, we adaptaded MagMax™ CORE Nucleic Acid Purificatioin Kit (ThermoFisher™) for a manual procedure using magnetic racks. Only reagents provided by the kit were used and the steps basically followed the automated ones. An initial 200 μl aliquot of each sample was mixed with 30 μl of beads mix (20 μl of magnetic beads and 10 μl of proteinase K) by pippeting up and down ten times, 700 μl of lysis-binding mix were then add and tubes were homogenized by vortexing, transferred to the magnectic rack where stayed for 1 min before supernatant removal. Five hundred microliters of Wash 1 were add to the samples, tubes were vortexed for 1 min, spinned for 10 sec at 500 rpm and then placed in the magnetic rack again for 1 min until supernatant removal, Wash 2 were used as described for Wash 1 and after supernatant removal the tubes stayed in rack with lids open to dry for 5 min. Elution buffer (90 μl) was then added to the tubes and they were homogenized in vortex for 3 min, spinned 10 sec and placed in the rack for 3 min until elute transfer to a new tube.

### Detection by quantitative Reverse Transcriptase Polymerase Chain Reaction (qRT-PCR)

Detection of SARS-CoV-2 were done with the set of primers and probe of the Charité protocol for amplification of E gene with some modifications that will be described here [13]. Reactions were done in a total volume of 20 μl, being 10 μl of 2X RT-PCR Buffer and 0,8 μl of Enzyme Mix of the AgPath-ID™ One-Step RT-PCR Reagents (ThermoFisher™), 0,8 μl of each primer and 0,4 μl of probe for E gene [13] both in a 5 μM concentration, 2,2 μl of nucleasse-free water and 5 μl of RNA sample. Amplification cycle was also adapted, starting in 50°C during 15 min for reverse transcriptase, followed by denaturation in 95°C for 10 min and 40 cycles in 95°C for 15 sec and 60°C for 45 sec.

For RNAse P amplification the set of primers and probes N1 and N2 of CDC diagnostic panel [14] were used in the following conditions: 20 μl total volume being 10 μl of 2X RT-PCR Buffer and 0,8 μl of Enzyme Mix of the AgPath-ID™ One-Step RT-PCR Reagents (ThermoFisher™), 1,5 μl of primers + probe, 2,7 μl of nuclease-free water and 5 μl of RNA sample. Amplification cycle for RNAse P detection was the same described above for the E gene.

## RESULTS

The method with the best performance, lower CT values to both SARS-CoV-2 or RNAse P and greater detection sensitivity of the control dilutions was the magnetic beads method MagMax™ that succeed to detect until a dilution of 10^−8^ of the control with a CT value of 38,47, all CT values to SARS-CoV-2 detection are listed in the Table 1, RNAseP detection results could be checked in the Table 2. MagMax™ and Biopur^®^ presented the lowest CT values and TRIzol™ performed slightly better than EasyExtract.

**Table 1:** Cycle threshould values for E gene detection of SARS-CoV-2 by different extraction methods

| ID | MagMax | TRIzol | Biopur | EasyExtract |
| --- | --- | --- | --- | --- |
| PC pure | 12,99 | 8,61 | 8,71 | ND |
| PC -1 | 14,01 | 14,35 | 15,86 | 24,10 |
| PC -2 | 17,53 | 21,54 | 23,05 | 22,76 |
| PC -3 | 20,86 | 25,82 | 26,40 | 25,91 |
| PC -4 | 24,43 | 28,00 | 27,98 | 28,90 |
| PC -5 | 27,74 | 32,90 | 31,83 | 32,96 |
| PC -6 | 30,99 | 36,21 | 34,53 | 35,97 |
| PC -7 | 34,26 | 37,39 | 37,95 | 39,14 |
| PC -8 | 38,47 | ND | ND | 37,36 |
| 6539 | 29,66 | ND | 35,90 | ND |
| 6499 | 33,25 | 32,55 | 33,03 | ND |
| 6842 | 33,69 | 35,38 | 34,97 | ND |
| 6321 | ND | ND | ND | ND |
| 6754 | 25,53 | 24,11 | 24,20 | 30,14 |
| 6928 | 28,75 | 29,08 | 29,67 | 35,57 |
| 6935 | 18,80 | 18,32 | 18,67 | 24,14 |
| 6938 | 20,26 | 20,11 | 18,51 | 27,33 |
| 6941 | 20,54 | 22,40 | 20,36 | 26,53 |

ID: final sample identification; PC: positive control; ND: not detected.

**Table 2:** Cycle threshould values for RNAse P detection of clinical samples by different extraction methods

| ID | MagMax | TRIzol | Biopur | EasyExtract |
| --- | --- | --- | --- | --- |
| 6539 | 19,29 | ND | 16,25 | 24,43 |
| 6499 | 24,71 | 30,41 | 23,86 | 29,38 |
| 6842 | 26,96 | 31,32 | 27,60 | 32,04 |
| 6321 | 25,87 | 33,6 | 25,90 | 31,73 |
| 6754 | 26,24 | 30,97 | 24,14 | 31,17 |
| 6928 | 28,41 | 30,68 | 27,69 | 33,61 |
| 6935 | 27,19 | 30,20 | 24,67 | 32,63 |
| 6938 | 23,66 | 27,16 | 21,67 | 29,09 |
| 6941 | 26,66 | 32,44 | 26,30 | 31,48 |

ID: sample identification; ND: not detected

Two singularities must be mentioned, bronchoalveolar lavage sample 3539 was negative in TRIzol™ technique probably due to it’s great viscosity, although it was initially diluted in MEM, which probably impaired technique execution mainly in the step of phase separation with chloroform. So, in this situation a greater dilution of the sample may be required to do this technique. When EasyExtract technique was done with the pure samples there was interference in qRT-PCR detection in control samples due to MEM coloration and in the clinical samples due to presence of inhibitors, taking into account that this method does not have a wash step. A second test was take with all the samples diluted in a 1:10 proportion, what resulted in the values utilized for comparison with the other methods and that are shown in Table 1 and 2, despite of one more thawn was done with the samples.

Concerning to the mean delay in CT values when compared to MagMax™, TRIzol™, Biopur^®^ and EasyExtract presented 0,39, 0,95 and 5,23 respectively. High delay in CT values of EasyExtract may impair detection in samples with low viral titer resulting in false negative results. Phenol-chloroform and silica column separation methods besides have presented few delay when compared with MagMax™ also presented similar values among itselves with 0,99 correlation.

Ultimately, agreement between positive and negative results of different methods when compared with the one with better performance MagMax™ was 94,44% for silica column separation method (Biopur^®^), followed by 88,89% for phenol-chroloform method (TRIzol™) and 77,78% for EasyExtract. Based on these results, the better alternatives of manual extraction methods to reduce chances of false negative results in Covid-19 diagnostics are magnetic beads extraction method MagMax™ and the silica column separation method (Biopur^®^).

## DISCUSSION

The sudden start of the current pandemic of a new coronavirus, now known as SARS-CoV-2 and rapid growth in number of suspects cases took scientists all over the world to run for a sensitive, especific and reliable diagnostic method to test the growing number of victims. After the first genomes sequences were published by chinese researchers it was possible to establish a qRT-PCR diagnostic protocol to large scale production and distribution [13, 14].

With the great demand of materials and reagents some laboratories face lack of products and/or long delivery time while a growing number of patients await for a diagnostic result. Nucleic acids isolation and purification is a major step in the diagnostic of Covid-19 and althogh the main laboratories have this proccess automated, many others around the world mainly in developing countries does not have this option. Besides of that, the already mentioned lack of products may lead automated laboratories to swift to manual proccess to accomplish the diagnostic requiriments.

In order to evaluate performance of alternative methods to the automated ones, in this study we have compared four different manual nucleic acids extraction methods for gene E SARS-CoV-2 and RNAse P detection. Being them the magnetic beads-based method MagMax™ CORE Nucleic Acid Purificatioin Kit (ThermoFisher™), column-based method Mini Spin Plus Kit (Biopur^®^), TRIzol™ (Invitrogen™) a phenol-chloroform based-method and the sample preparation EasyExtract.

Unfortunately, besides being atractive due to fastness and low handling the EasyExtract method presented pour results, not being reliable mainly in clinical samples with low viral titers. Also, due to absense of a wash step this technique maintains the presence of inhibitors and a sample dilution may be required. Still, in emergency situations it can be used as a last resort.

Phenol-chloroform-based method evaluated here presented satisfactory results, this technique possess the advantage of allowing RNA isolation and having protective characteristics for that. Although the agreement of positive-negative results were of only 88,89% comparing to MagMax™, it presented the lowest CT delay among the other techniques (0,39). Lastly, this method presents some cons given that possess a laborious and time consuming process and requires the use of toxic, harmful and irritating reagents. With this in mind, it is still a good alternative when necessary.

In this study, the silica column-based method presented results almost as good as the magnetic beads method taking into account that have a delay of 0,95 and a positive-negative agreement of 94,44%, proving itself as a good manual kit option. Both techniques have also a similar process time.

Similar to our findings, another study that have compared the magnetic beads with silica column-based methods for human gut microbial community profiling obtained higher amounts of nucleic acids extracted with the magnetic beads method and greater species diversity, it also showed that manual extraction presented similar results to the automated ones when taking into account the reproducibility of microbial profiles, althogh it might be different for nucleic acids yields [15]. However, most studies that have done comparison of different extraction methods have used feces samples to bacteria detection [16, 17].

## CONCLUSION

In this work we have compared four different types of manual extraction methods for SARS-CoV-2 detection by qRT-PCR with clinical samples and control dilutions. We aimed to evaluate how reliable each method is for diagnostic purposes and to propose alternatives when usual methods are not available. In this regard, magnectic beads and silica column-based methods are convenient and reliable choices, phenol-chloroform-based method could also be chosen as a alternative method unlike to EasyExtract that is not a trustworthy option.

## ACKNOWLEDGMENTS

We would like to thank to Coordenação de Aperfeiçoamento de Pessoal de Nível Superior – CAPES for the graduate scholarships, Financiadora de Estudos e Projetos – Finep and RedeVírus Ministério da Ciência, Tecnologia, Inovações e Comunicações – MCTIC for the research funding.

## Notes

### Competing Interest Statement

The authors have declared no competing interest.

